# Transfer of weight information depends differently on used hand and handedness for perception and action

**DOI:** 10.1101/2022.04.25.489365

**Authors:** Vonne van Polanen

## Abstract

When lifting an object sequentially with the two hands, information about object weight can be transferred from one hand to the other. This information can be used to predictively scale fingertip forces and to form a perceptual estimation about the object’s weight. This study investigated how weight information can be transferred between the two hands and how this depends on the used hand and handedness of the participant. Right- and left-handed participants lifted light and heavy objects with the right or left hand in a pseudorandomized order and estimated the object’s weight. Results showed that predictive force scaling depended on the previously lifted object, with higher forces rates if a previous object was heavy. This known effect of sensorimotor memory was mostly independent of the used hand and handedness, indicating that weight information could be transferred for fingertip force scaling. Furthermore, a perceptual bias depending on the previous lifted object was found, with lower weight estimations when the previous object was heavy compared to light. In contrast to the results on force scaling, a transfer of this perceptual bias was only found in left-handed participants. In addition, right-handers only showed a bias when they used their dominant hand. These findings indicate that the transfer of weight information depends differently on the used hand and handedness for perceptual estimations and predictive force scaling.

## 1 INTRODUCTION

For skilful object lifting, fingertip forces must be predictively scaled towards the weight of the object. Predictive scaling is important, because feedback processes are often too slow and will result in less smooth movements [1, 2]. When the weight of the object is unknown, forces are typically based on the weight of the previously lifted object, an effect that has been termed sensorimotor memory [2-6]. Specifically, when a previous object was light, the force rates for the current lift are lower than if a previous object was heavy.

Previous studies showed that this sensorimotor memory can be maintained for hours [7, 8] and the stored information can also be transferred between hands [9-12]. That is, when an object is lifted with one hand, the sensorimotor experience is used to scale forces for lifting an object with the other hand. The transfer of the sensorimotor memory for force scaling has generally been found not to depend on the used hand (i.e. right to left or left to right hand) and was equal in both directions [9, 12], although one study found larger effects when transferring from the right to the left hand in right-handed participants [11]. Furthermore, a more efficient transfer was found in right-handed compared to left-handed participants [12]. Differences between hands and handedness could suggest that processing of sensory information is lateralized to one hemisphere. However, since few studies on object lifting compared whether effects depended on the used hand or handedness, results on laterality and handedness remain scarce.

When lifting an object, the acquired sensory information can also be used to provide a perceptual judgement of the object’s weight. However, this estimate is not always veridical. Previous studies have shown that a previous lifted object can affect the estimated weight of the following lifted object [5, 6, 13]. When the lifted object is preceded by a heavy object, it is judged to be lighter than when preceded by a light object, indicating a perceptual bias. The similar effect of previous lifts on force scaling and weight perception suggests that they originate from the same mechanism. Since no perceptual bias was found when objects were placed on the hand instead of being lifted, it has been suggested to originate from the corrections to the fingertip forces [5]. In support of this hypothesis, this study found a correlation between force and perceptual parameters. However, in a follow-up study, this correlation was not found [6], resulting in conflicting findings on the relation between force control and weight perception based on previous lifts.

If the sensory information about object weight used for force scaling and weight perception stem from the same mechanism, it would be expected that this information is transferred in a similar way for both processes as well. In contrast to the transfer of weight-related information for force scaling, less is known about the transfer of information for weight perception. A transfer of the perceptual bias, although weaker, was found from the left to the right hand [13], but this study was limited to a single trial. More research on the transfer of perceived weight has been performed in the context of the size-weight illusion. In this illusion, a small object is perceived to be heavier than a large object despite having the same mass [14]. It has been found that the illusion effect can be transferred from one hand to the other, although this effect was only visible with a transfer from the left to the right hand and not in the other direction [9]. However, this illusion has been shown to be independent from force scaling when looking at behaviour after repeated lifts [15, 16].

All in all, it remains unclear whether perceptual information about object weight can be transferred across hands and how this depends on the used hand and handedness of the participant. To address this issue, an experiment was performed where participants lifted objects of different weights. The perceptual bias based on a previous lifted object was used to examine whether weight information can be transferred from one hand to the other. Participants either lifted objects sequentially with the same hand or with different hands to access the ability to transfer sensory information. Force scaling and weight estimations were measured to investigate the sensorimotor memory and the perceptual bias of previously lifted objects. In addition, it was measured whether the perceptual bias depended on the direction of transfer (from left to right, or from right to left) and on handedness (right-handed and left-handed participants).

This study will also provide insight into the potential relation between the sensorimotor memory and perceptual biases from previous experience. If both processes stem from the same mechanism, it is expected they will depend similarly on the used hand and handedness and that effects correlate with each other. Since the sensorimotor memory can be transferred across hands [9-12] and the perceptual bias was present in a transfer from the left to the right hand [13], it was expected that the perceptual bias would be transferred between hands as well.

## 2 METHODS

### 2.1 Participants

Sixty participants were recruited for this study and divided over four groups. Handedness was assessed with the Edinburgh Handedness Inventory [17]. Twenty right-handed (mean LQ=86) participants were assigned to the right-hand group (22.1 years, age range 18-27 years, 11 females) and another twenty right-handed participants (mean LQ=88) were assigned to the left-hand group (21.8 years, 18-26 years, 9 females). Similarly, ten left-handed participants were placed in a right-hand (mean LQ=–85, 23.5 years, age range 19-31 years, 5 females) and another ten in a left-hand group (mean LQ=– 86, 22.2 years, age range 20-27 years, 6 females). All participants provided written informed consent before participating. Ethical approval was obtained from the local ethical committee of KU Leuven.

### 2.2 Apparatus

Two force sensors (Nano 17, ATI Industrial Automation) were attached to a manipulandum in which differently weighted objects could be placed (Fig. 1). The force sensors were covered with sandpaper (No. P600) to increase the friction of the surface. Forces were sampled in three directions with a sample frequency of 1000 Hz.

**Fig. 1.**
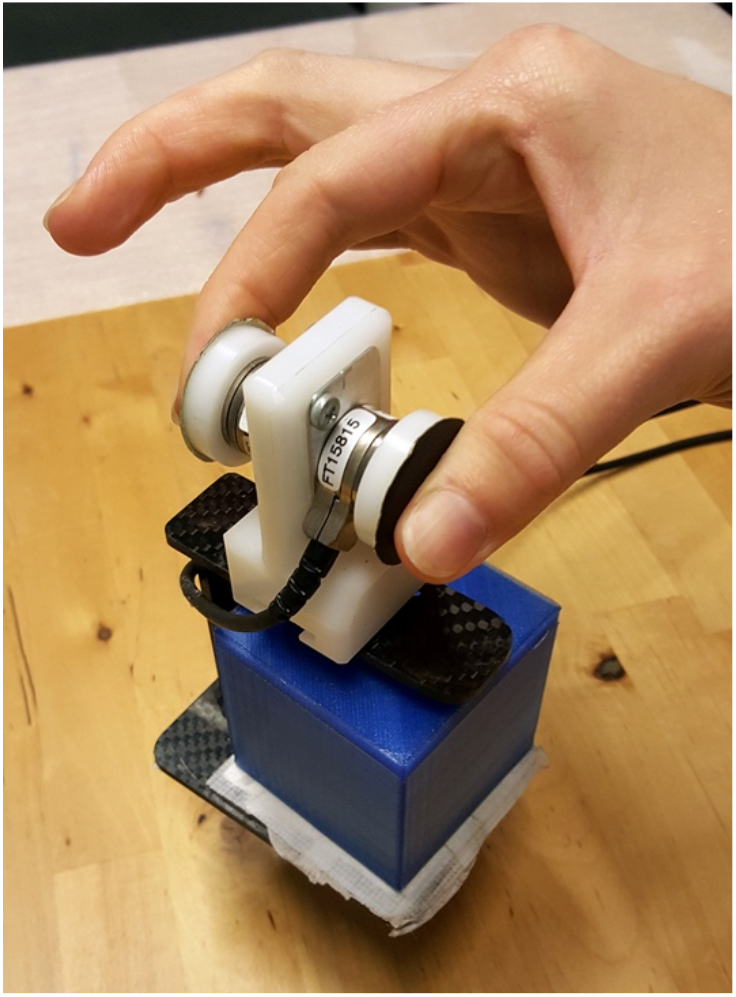
Experimental manipulandum with force sensors, covered with sandpaper. Different cubes could be placed in the manipulandum.

Three 3D-printed objects were used in the experiment, which all measured 5 × 5 × 5 cm. They were filled with different amounts of lead shot to created weights of 105, 317 and 525 g. A fourth object of 260 g was used for practice trials. The manipulandum, including de force sensors, weighted approximately 120 g. Therefore, the lifted weights were 2.2, 4.3, 6.3 and 3.8 N, respectively. The cubes were placed under a paper cover to prevent visual cues.

To prevent participants seeing the experimenter changing the objects, participants wore PLATO glasses (Translucent Technologies) or a switchable screen (Magic Glass) was placed in front of them. The glasses and the screen could switch between an opaque and transparent state.

### 2.3 Experimental Procedure

Participants were seated at a table on which the manipulandum was placed and rested both hands on the table. They received an auditory cue indicating which hand should be used for the next trial. This cue was a computer voice saying ‘left’ or ‘right’. 1.5 s after this cue, the glasses or screen became transparent and participants reached for the manipulandum and lifted it with their thumb and index finger placed on the force sensors. They were told to lift the manipulandum straight, at a comfortable speed and to a height of ±5 cm and hold it steady. When the glasses or screen turned opaque again after 3 s, they replaced the manipulandum on the table again. Next, they were asked to estimate the weight of the lifted object on a self-chosen scale with no predetermined lower or upper limit (magnitude estimation). The experimenter then changed the object for the next trial. Before the experiment, 5 trials for each hand in a randomized order were performed with the practice object of 260 g to get familiar with the task procedure.

Trials were presented in a pseudo-randomized order where the previous and the current lift were carefully controlled. There were four weight conditions where the order of the light (105 g) and heavy (525 g) objects were varied: light-light (LL), heavy-light (HL), light-heavy (LH) and heavy-heavy (HH). Here, the first weight refers to the object weight on the previous lift, and the second weight refers to the object weight of the current lift. In addition, the hand used in the previous and current lift was varied as well: either it was the same hand (S) or a different hand (D). Therefore, there were 8 conditions in the experiment: LLS, HLS, LHS, HHS, LLD, HLD, LHD, HHD. In the conditions where a different hand was used, the transfer of information from one hand to the other could be assessed. The used hand in the current lift and the previous lift differed across the four groups. The two groups (right-handed and left-handed participants) in the right-hand groups always performed the current lift with their right hand. Therefore, they would lift objects in the same hand conditions twice with their right hand, whereas in the different hand conditions they would lift the previous object with their left hand and the current one with their right hand. In contrast, the two groups in the left-hand groups performed the current lift with their left hand. For those groups, the same hand conditions consisted of two lifts with the left hand and the different hand conditions were lifts first with the right hand followed by the left hand. In this way, the transfer towards the right hand and towards the left hand could be examined in the right-hand and left-hand group, respectively. An overview of the conditions for each participant group is shown in Table 1.

**Table 1.**
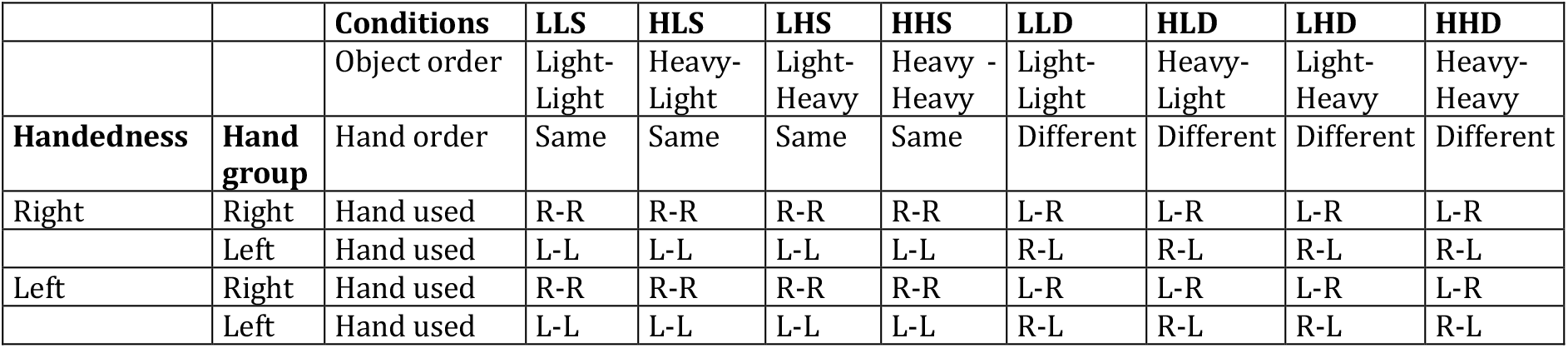
Overview of the participant groups and conditions. For each condition, the object weight in the previous and current trial (object order) is indicated (Light or Heavy). The used hands in the previous and current trial (hand order), where either the same (S) or different (D) hands are used, could be the left (L) or right (R) hand, dependent on the hand group.

Each of the 8 conditions were presented 10 times, giving a total of 80 trials. To provide more variation in the weights used in the experiment, a third object of 317 g was used as a dummy object. This object was lifted in about 10% of the trials (15 trials), either with the left or right hand. These trials, and trials following dummy trials were not analysed. In addition, the sequence could also contain trials with the wrong hand order (e.g. left hand followed by right hand for the left-hand group; max 40 trials, this varied from sequence to sequence), which were also not analysed. Finally, the first trial was not preceded by any lift, so it was not analysed either. A total of 151 trials were presented of which at least 80 were entered into the analysis. The trial sequence was different for each participant.

### 2.4 Data analysis

Participants’ weight estimations were normalized to z-scores. Force data were filtered with a second order low-pass bidirectional Butterworth filter, with a cut-off frequency of 15 Hz. Load forces (LF) were the sum of the vertical forces and grip forces (GF) the average of the horizontal forces. These forces were differentiated with respect to time to obtain load forces rates (LFR) and grip force rates (GFR). From the force rates, the first peaks of the forces rates were of interest, since these reflect the planned force [1]. The first peaks (peakLFR1 and peakGFR1) were calculated between the time points of contact (GF>0.1 N) and just after lift-off (50 ms after lift-off, where LF equals object weight). To exclude peaks that were noise or accidental bumps when contacting the manipulandum, the first peak had to be at least 30% of the maximum force peak as measured between these time points [6]. Finally, the loading phase duration (LPD) was calculated as the time between onset of LF (LF>0.1 N) and lift-off (LF>object weight).

Trials in which the wrong object was placed, the wrong hand was used or the object was not lifted were removed from the perceptual analysis, as well as trials following these erroneous trials since the order would be wrong (total per group: 12, 0.7%, in right-handed with right hand; 8, 0.5%, in right-handed with left hand; 4, 0.5%, in left-handed with right hand; 3, 0.4%, for left-handed with left hand). In addition, trials in which the object was lifted twice, too early, or pushed before lifting, were also removed from the force analysis (total per group: 19, 1.2%, in right-handed with right hand; 19, 1.1%, in right-handed with left hand; 8, 1%, in left-handed with right hand; 6, 0.7%, for left-handed with left hand).

### 2.5 Statistics

Statistical analyses were performed with SPSS Statistics (IBM, version 28). The parameters of interest (perceptual answers, peakGFR1, peakLFR1, LPD) were averaged for each condition and in each group. First, it was examined whether any differences between the groups were present. Therefore, a mixed analysis of variance (ANOVA) was performed with the following between factors: handedness (2 levels: left-handed, right-handed), current hand (2 levels: left, right) and the following within factors: previous hand (2 levels: same, different), previous weight (2 levels: light, heavy) and current weight (2 levels: light, heavy). If there were significant effects of both between factors and within factors, separate repeated measures ANOVAs were performed for each group with the 3 within factors. Further post-hoc tests were performed with paired samples t-tests with a Bonferroni corrections. An alpha level of 0.05 was used.

Finally, it was tested whether the perceptual biases were related to the effects on the force parameters. The differences with respect to previous weight were used to investigate the relation between effects on perception and force. Similar to the z-scored perceptual answers, the peakGFR1 and peakLFR1 were converted to z-scores and the difference between a previous light and a previous heavy weight was calculated. The difference for the perceptual answers reflects the perceptual bias and the difference for the force parameter reflects the sensorimotor memory effect. A mixed model was used with the peakGFR1 or the peakLFR1 difference as predictor, in two separate analyses. The perceptual difference was the dependent variable and the peakGFR1 or peakLFR1 difference was entered as a covariate in a first-order autoregression model with maximum likelihood estimation. The same factors as in the mixed ANOVA (handedness, current hand, previous hand and current weight) were used in the model.

To test whether the parameters were related in general on a trial-by-trial basis, a similar mixed model was used where all trials of participants were entered. The trials were compared within participants, regardless of current used hand or previous weight. The peakGFR1 or peakLFR1 were entered as covariates, the perceptual answer as dependent variable and a first-order autoregression model with maximum likelihood estimation was used. As fixed factors, handedness and current hand were used, to distinguish between participant groups. This analysis was performed for current light and current heavy objects separately.

## 3 RESULTS

In this study, participants lifted light and heavy objects and estimated their weight. It was investigated how participants planned their fingertip forces and estimated the weight of the current lift depending on the previous object they lifted. Participants performed the current lift with their right or left hand and were either right- or left-handed, resulting in four participants groups. Furthermore, the current lift was either performed with the same hand as the previous lift, or with a different hand to investigate whether information from the previous lift could be transferred to the other hand. All data used for analysis and figures and analysis scripts can be found on https://osf.io/kgrez/.

### 3.1 Perceptual biases depend on the used hand and handedness

Participants estimated the weight of each object after they had lifted it. First, it was investigated whether the estimates were different depending on the different participant groups. The mixed ANOVA indicated significant effects of the between factors handedness (F(1,56)=5.8, p=0.020, η_p_^2^=0.09) and current hand (F(1,56)=7.5, p=0.008, η_p_^2^=0.12), as well as an interaction between current weight × previous weight × handedness (F(1,56)=8.7, p=0.005, η_p_^2^=0.13). In addition, there were also effects of the within factors current weight (F(1,56)=21989.0, p<0.001, η_p_^2^=1.00) and previous weight (F(1,56)=23.8, p<0.001, η_p_^2^=0.30) and interactions of previous hand × previous weight (F(1,56)=10.1, p=0.002, η_p_^2^=0.15) and current weight × previous weight (F(1,56)=15.4, p<0.001, η_p_^2^=0.22). Because of the significant between factors, separate repeated measures ANOVAs were performed for each participant group. Especially the effect of previous weight was of interest, which would indicate the presence of a perceptual bias. Results for the perceptual estimates are shown in Fig. 2.

**Fig. 2.**
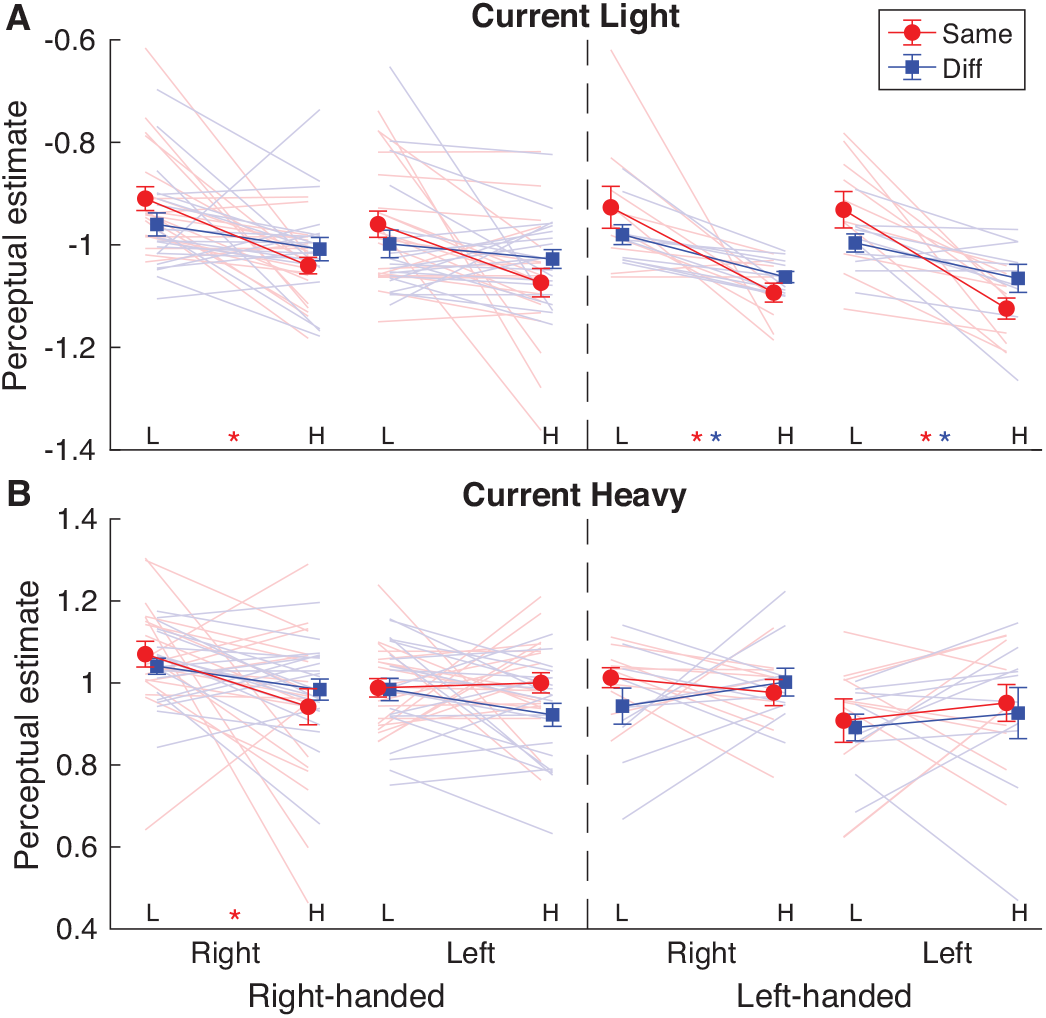
Results for the perceptual estimates in each condition for current light (**A**) and current heavy (**B**) objects. Lines are shown from a previous light (L) to a previous heavy (H) lift for each participant group, who were right-handed (before dashed line) or left-handed (after dashed line) and used their right or left hand in the current lift. In the previous lift, the same hand (red, Same) or a different hand (blue, Diff) was used. Solid lines and error bars represent means and standard errors. Light shaded lines indicate individual participants. Asterisks indicate a significant difference between a previous light and previous heavy object, representing a perceptual bias (red: Same, blue: Different hand condition).

#### 3.1.1 Right-handed with right hand

The 2 (previous hand) × 2 (current weight) × 2 (previous weight) ANOVA showed an effect of previous weight (F(1,19)=17.9, p<0.001, η_p_^2^=0.49). This effect showed that there was a perceptual bias, where an object was judged to be heavier if it was preceded by a light object, compared to a heavy object. However, there was also an interaction between previous hand × previous weight (F(1,19)=7.3, p=0.014, η_p_^2^=0.49=0.28) that indicated that the perceptual bias was only seen when the objects were lifted with the same hand (p<0.001), not with different hands (p=0.120). Furthermore, a main effect of current weight was found (F(1,19)=9592.2, p<0.001, η_p_^2^=1.00). As expected, heavy objects were estimated as being heavier than light objects.

#### 3.1.2 Right-handed with left hand

When the currently used hand was the non-dominant hand of right-handed participants, there were effects of current weight (F(1,19)=6544.4, p<0.001, η_p_^2^=1.00) and previous weight (F(1,19)=4.8, p=0.041, η_p_^2^=0.20). There was also a 3-way interaction of previous hand × current weight × previous weight (F(1,19)=6.4, p=0.020, η_p_^2^=0.25). The main effect of previous weight indicated that when the previous object was light, perceptual ratings were higher than when the previous object was heavy. However, none of the post-hoc tests in the 3-way interaction showed a significant effect of previous weight in any condition (all p>0.07). Neither were there any significant differences between conditions in which the same or different hands were used (all p>0.21). Unsurprisingly, the heavy object was judged to be heavier compared to the light object, which was the case in all conditions (all p<0.001).

#### 3.1.3 Left-handed with right hand

For left-handers when using the right hand, there was also an effect of previous weight (F(1,9)=5.3, p=0.047, η_p_^2^=0.37), indicating that when the previous object was light, a higher estimate was given than when the previous object was heavy. The interaction between current weight × previous weight (F(1,9)=6.8, p=0.028, η_p_^2^=0.43) indicated that this perceptual bias was only seen when the current object was light (p=0.022), not heavy (p=1.00). Finally, a main effect of current weight (F(1,9)=8018.7, p<0.001, η_p_^2^=1.00) showed that participants judged the object as being heavier when it was heavy compared to light, regardless of the previous weight (both p<0.001).

#### 3.1.4 Left-handed with left hand

When left-handers were using the left hand, there was an interaction between current weight and previous weight (F(1,9)=18.6, p=0.002, η_p_^2^=0.67). When the previous object was heavy, a lower weight estimate was given than when the previous object was light, but only when the current object was light (p=0.002), not heavy (p=1.00). A main effect of current weight was found (F(1,9)=3348.7, p<0.001, η_p_^2^=1.00), where again the heavy object was perceived to be heavier than the light, regardless of the previous weight (both p<0.001).

Overall, participants could differentiate between light and heavy objects in all conditions. There was a perceptual bias due to object order, but this depended on the handedness, hand used and the weight of the object (see Fig. 2). For right-handers, only a perceptual bias was found when using the same, dominant hand. Therefore, no transfer or perceptual information was found for right-handed participants. For left-handed participants, the perceptual bias did not seem to depend on which hand was used, nor whether same or different hands were used, but it was only seen when lifting light, not heavy objects.

### 3.2 Peak grip force rates: effects of previous object depend on the used hand

To examine the force scaling for the lifted object, the first peak of the force rate was calculated. Next, the different participant groups were compared with a mixed ANOVA. For the peakGFR1, a significant interaction with the two between factors previous hand × current hand (F(1,56)=5.4, p=0.024, η_p_^2^=0.09) was found, as well as a 3-way interaction between current weight × handedness × current hand (F(1,56)=5.6, p=0.021, η_p_^2^=0.09). The mixed ANOVA also revealed main within factor effects of current weight (F(1,56)=20.1, p<0.001, η_p_^2^=0.270) and previous weight (F(1,56)=158.1, p<0.001, η_p_^2^=0.74), and an interaction between previous hand × previous weight (F(1,56)=31.0, p<0.001, η_p_^2^=0.36). Since the significant between factors indicate differences between the participant groups, separate ANOVAs were performed for each participant group. The peakGFR1 results are illustrated in Fig. 3.

**Fig. 3.**
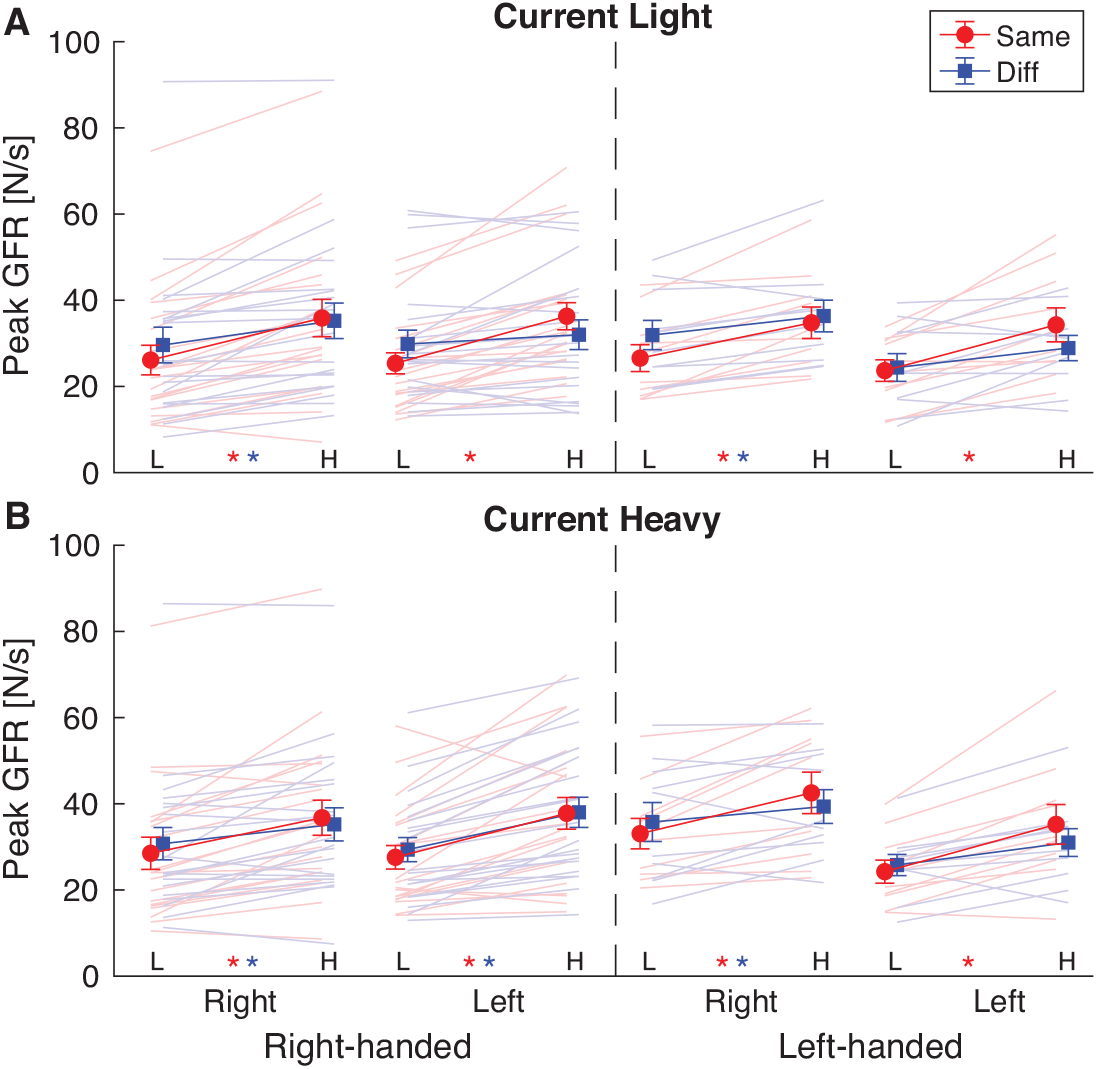
Results for the first peak grip force rate (peak GFR) in each condition, for current light (**A**) and current heavy (**B**) objects. Lines are shown from a previous light (L) to a previous heavy (H) lift for each participant group, who were right-handed (before dashed line) or left-handed (after dashed line) and used their right or left hand in the current lift. In the previous lift, the same hand (red, Same) or a different hand (blue, Diff) was used. Solid lines and error bars represent means and standard errors. Light shaded lines indicate individual participants. Asterisks indicate a significant difference between a previous light and previous heavy object, representing different force scaling (red: Same, blue: Different hand condition).

#### 3.2.1 Right-handed with right hand

A repeated measures ANOVA was performed for the group of right-handed participants that used their right hand in current lifts. The peakGFR1 was larger when the previous object was heavy compared to when it was light (previous weight: F(1,19)=66.7, p<0.001, η_p_^2^=0.78). There was also an interaction of previous hand × previous weight (F(1,19)=11.2, p=0.003, η_p_^2^=0.37). Both when using the same hand (p<0.001) or different hands (p<0.001), a previous heavy object resulted in higher force rates than a previous light object. When the previous object was light, a higher force rate was used when different hands were used compared to using the same hands (p=0.012), while there was no difference for a previous heavy object (p=0.73).

#### 3.2.2 Right-handed with left hand

When right-handers were using the left hand, there were effects of current weight (F(1,19)=6.9, p=0.016, η_p_^2^=0.27) and previous weight (F(1,19)=51.4, p<0.001, η_p_^2^=0.73). However, since there were also interactions of previous hand × previous weight (F(1,19)=11.3, p=0.003, η_p_^2^=0.37), current weight × previous weight (F(1,19)=7.3, p=0.014, η_p_^2^=0.28) and a three-way interaction between previous hand × previous weight × current weight (F(1,19)=6.0, p=0.024, η_p_^2^=0.24), the effects should be interpreted in light of these interactions. Participants scaled their forces differently depending on the previous lift. When a previous object was heavy, the peakGFR1 was higher than when the previous object was light, which was the case when using the same hands for current light (p<0.001) and current heavy (p=0.003) objects. This was also the case when using different hands, but only when the current object was heavy (p<0.001), not light (p=1.00). Furthermore, there was only a difference between light and heavy objects when the previous object was heavy and different hands were used (HLD vs HHD, p<0.001), where the force rate was higher for heavy objects. When the current object was light and the previous object as well, peakGFR1 was lower when using the same hand compared to different hands (LLS vs LLD, p=0.02).

#### 3.2.3 Left-handed with right hand

For left-handers using the right hand also an effect of previous weight was found (F(1,9)=44.3, p<0.001, η_p_^2^=0.83), that showed that when the previous object was heavy, the peakGFR1 was higher than when the previous object was light. In addition, the peakGFR1 was higher when the current weight was heavy compared to light (effect current weight: F(1,9)=9.5, p=0.013, η_p_^2^=0.51). However, there was also an interaction between previous hand × current weight (F(1,9)=5.3, p=0.047, η_p_^2^=0.37), which indicated that this difference between object weights was only present when using the same hands (p=0.015), not when using different hands (p=0.45).

#### 3.2.4 Left-handed with left hand

When the left-handed group used the left hand in current lifts, there was an effect of previous weight (F(1,9)=31.5, p<0.001, η_p_^2^=0.78). When the previously lifted object was heavy, the peakGFR1 was higher than when the previous lift was light. There was also an interaction between previous hand and previous weight (F(1,9)=8.3, p=0.018, η_p_^2^=0.48). This indicated that the difference between a previous light and heavy object was only significant when using the same left hand (p=0.001), but just failed to reach significance when using different hands (p=0.052).

To summarize, the grip force rates depended on the previous lifted weight: a higher force rate was seen when the previous object was heavy compared to light. When using the same hand, this effect was visible in all groups, whether the left or right hand was used. However, while a transfer of grip force rate to the right hand was found in all participant groups, the transfer to the left hand was less pronounced. For the right-handed participants that used their left hand, a transfer was only seen for heavy objects, whereas in left-handed participants no transfer of grip force to the left hand was seen for both object weights.

### 3.3 Peak load force rates: effects of previous object are independent of used hand and handedness

For the peak load force rates, the participant groups were also compared in a mixed ANOVA. However, the peakLFR1 did not depend on the handedness or current used hand in the task. Therefore, the groups will not be discussed separately and are shown pooled in Fig 4A. The mixed ANOVA showed a significant effect of previous weight (F(1,56)=92.5, p<0.001, η_p_^2^=0.62), which indicated that the peakLFR1 was higher when the previous object was heavy compared to light. Furthermore, there were effects of current weight (F(1,56)=46.6, p<0.001, η_p_^2^=0.45) and of previous hand (F(1,56)=5.3, p=0.025, η_p_^2^=0.09). The main effect of current weight demonstrated that peakLFR1 was higher when the current object was heavy compared to light. When using the same hands, the force rate was higher than when using different hands. However, there was also an interaction between previous hand × previous weight (F(1,56)=11.5, p=0.001, η_p_^2^=0.17) which indicated that this hand effect was only seen when the previous object was heavy (p<0.001), not light (p=0.24).

**Fig. 4.**
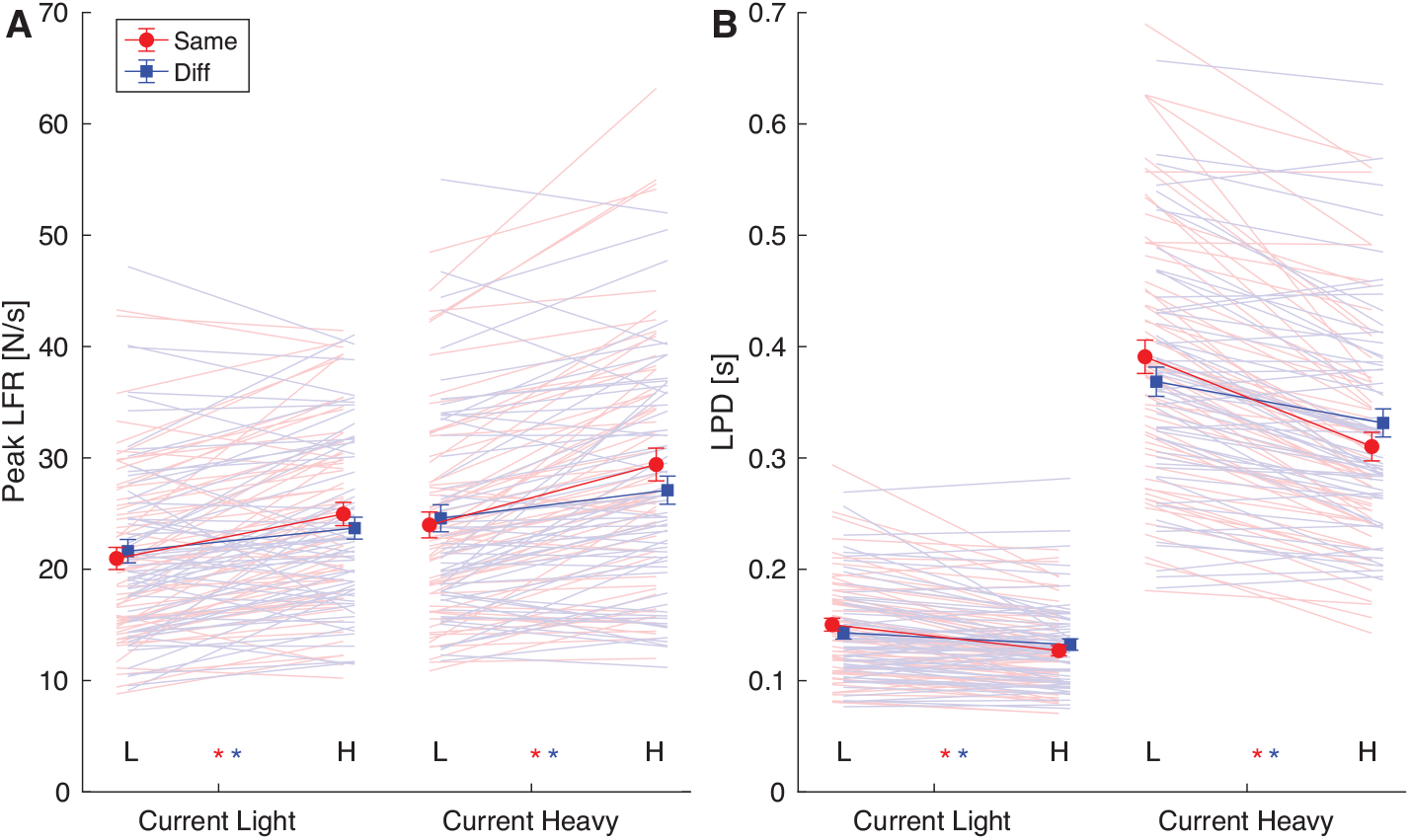
Results for the first peak load force rate (peak LFR, **A**) and loading phase duration (LPD, **B**). Lines are shown from a previous light (L) to a previous heavy (H) lift pooled over all participant groups (right- and left-handed, using right and left hand in current lift). Solid lines and error bars represent means and standard errors. Light shaded lines indicate individual participants. Asterisks indicate a significant difference between a previous light and previous heavy object, representing different force scaling (red: Same, blue: Different hand condition).

### 3.4 Loading phase duration: effects of previous object are independent of used hand and handedness

The LPD was not different between the different participant groups, as the mixed ANOVA showed no significant effects or interactions with the between factors. Therefore, results are further discussed pooled across all participants (Fig 4B). When the previous object was heavy, the LPD was shorter than when the previous object was light (previous weight: F(1,56)=207.2, p<0.001, η_p_^2^=0.79). There was also a significant effect of current weight (F(1,56)=507.2, p<0.001, η_p_^2^=0.90) that indicated that the LPD was longer when objects were heavy compared to light. In addition, significant interactions between previous hand × previous weight (F(1,56)=45.2, p<0.001, η_p_^2^=0.45), current weight × previous weight (F(1,56)=64.7, p<0.001, η_p_^2^=0.54) and a 3-way interaction of previous hand × current weight × previous weight (F(1,56)=10.0, p=0.003, η_p_^2^=0.15) were found. The difference between light and heavy objects, as well as differences between previous heavy and previous light objects were significant in all post-hoc tests (all p<0.02), revealing no new insights. The difference between using the same hand and different hands was only significant when the object was heavy (LH and HH conditions, both p<0.008), but not when the object was light (p>0.26). Here, the LPD was shorter when the same hands were used compared to different hands.

All in all, the peakLFR1 and the LPD depended on the weight of the current object and the previous object. There did not seem to be any differences between the hand that was used or the hand preference of the participant.

### 3.5 Correlations between perceptual biases and force parameters

The results described above show effects of a previous lifted object on perceptual answers (perceptual bias) and force scaling (sensorimotor memory). Individual participant data for all conditions are shown in Fig 5. as differences between a previous heavy and a previous light lift. Here, positive values on the y-axis (perception) indicate a perceptual bias in the expected direction and negative values on the x-axis (force scaling) indicate the expected effect from the sensorimotor memory. When values fall within the white square, both differences are in the expected direction. In line with the results as described above, this is mainly the case for lifting light objects and when lifting with the same hand.

**Fig. 5.**
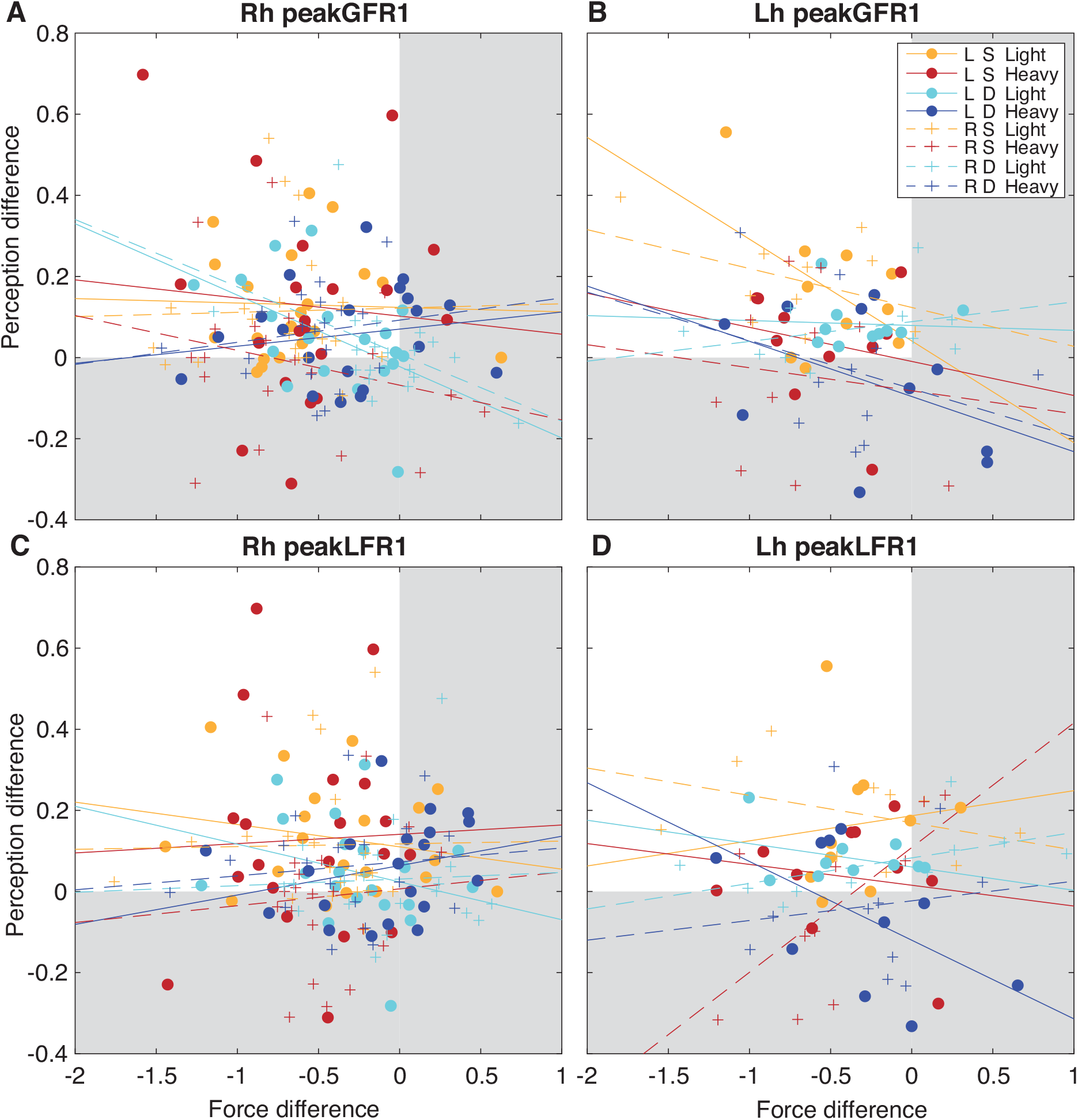
Correlations between perceptual bias and force scaling difference. The difference was calculated between a previous heavy and a previous light object, for the first peak of grip force rates (peakGFR1, **A-B**) and load force rates (peakLFR1, **C-D**). Data are shown for each participant group (**A**,**C** Rh=right-handed, **B**,**D** Lh=left-handed), and the use of the left hand (L, circles) or right hand (R, plusses). Colours indicate the use of the same (S) or different (D) for a current light or heavy object. Lines indicate the regression lines and points represent individual participants. Values within the white square indicate points in the expected direction (perceptual bias and sensorimotor memory). Note that none of the correlations were significant after Bonferroni correction.

To investigate the relation between effects of previous object on the perceptual answers and the force scaling, a mixed model was performed on the differences between a previous light and a previous heavy object. In the mixed model using the peakGFR1 as predictor, an effect of peakGFR1 was found (F(1,231)=8.3, p=0.004), in addition to an interaction of handedness × previous hand × current weight × peakGFR1 (F(1,223)=6.8, p=0.010). This suggest that the effects of the previous weight on the grip force rate seem to be related to the perceptual biases. Finally, a main effect of current weight was also found (F(1,231)=5.8, p=0.017), but this effect was already discussed for the ANOVA results.

For the peak load force rate, interactions of current hand × peakLFR1 (F(1,238)=4.4, p=0.038) and of handedness × current hand × current weight × peakLFR1 (F(1,235)=5.5, p=0.020) were found. This suggests that the peakLFR1 difference was related to the perceptual bias, although this depended on the specific condition. Finally, other effects of previous hand (F(1,201)=12.0, p=0.001), current weight (F(1,211)=6.9, p=0.009) and an interaction between handedness × current weight (F(1,211)=5.9, p=0.016) were found. Since effects of these factors were already discussed in the results of the ANOVA, they will not be further discussed here.

Since the peak force rates also interacted with some conditions, separate correlations were performed between the peak force differences and the perceptual biases in each condition (2 handedness × 2 current hand × 2 previous hand × 2 current weight = 16 correlations for peakGFR1 and for peakLFR1). The correlations are illustrated in Fig. 5. However, after Bonferroni correction, none of these correlations were significant (p>0.32).

For the trial-by-trial analysis, it was found that peakGFR1 was a significant predictor for the perceptual estimate, both for light (F(1,2447)=55.0, p<.001) and heavy (F(1,2477)=5.3, p=0.022) objects. PeakLFR1 was only a significant predictor in light (F(1,2436)=30.9, p<.001), but not for heavy objects (F(1,2476)=2.9, p=0.086). This suggests that on a trial-by-trial basis, perceptual estimates were related to the grip force rate peak and also to the load force rate peak, but for the latter parameter only when lifting light objects.

## 4 DISCUSSION

The main aim of this study was to investigate how weight information can be transferred between the hands when lifting objects. Specifically, the effects of the used hand and handedness on force scaling and the perceptual bias that occurs in weight perception depending on the previously lifted object were examined. Four groups of participants who were either right- or left-handed lifted objects of different weights and used their left or right hand to lift these objects. They sequentially lifted two objects with the same hand or with different hands and estimated the objects’ weight. It was found that the previously lifted weight affected anticipatory force scaling and that information from the previous lifted weight could be transferred across hands for predictive force scaling. However, while a perceptual bias was found when using the same hands, particularly for light objects, this information could only be transferred in left-handed but not in right-handed participants.

When an object was lifted after a light object, it was perceived to be heavier than when it was lifted after a heavy object, indicating that a perceptual bias was present. For right-handed participants, this bias was only present when they lifted the objects with the same hand, not with different hands. In contrast, for left-handed participants, the bias was present irrespective of the hand that was used. This indicates that the weight information could be transferred between hands for the left-handers and this was the case both when transferring from the right to the left hand and from the left to right hand. A possible explanation for this handedness effect could be a better transfer of information due to a stronger connection between hemispheres, since the corpus callosum is smaller for strong right-handers compared to left and mixed-handed participants [18]. Previous studies on motor skill learning have shown similar advantages for left-handed compared to right-handed participants. Skill transfer was only seen in left-handers, not right-handers [19], or was superior for left-handers [20].

Besides the differences in transfer of information, also when using the same hand, differences with respect to handedness were seen. The perceptual bias was only present when right-handed participants used their right, dominant, hand, while a bias was seen irrespective of the used hand for left-handed participants. While there are few studies on the effect of handedness on weight perception, no apparent differences in weight discrimination are known between right- and left-handers [21-23], but the size-weight illusion seems to differ between hands for right-handed and not for left-handed participants [12]. Furthermore, weight discrimination is better for the dominant than the non-dominant hand [21], which would explain why a bias was only found for the dominant hand in the right-handers. In motor skill learning it has been found that the difference between the dominant and the non-dominant hand is less apparent in left-handed compared to right-handed participants [24-26] and also depends on the strength of handedness [20]. Since being left-handed is less common than being right-handed, environmental circumstances could influence the degree to which left-handers use their dominant and non-dominant hand, because they have to adapt to a world that is designed for right-handers. Hence, they might be more accustomed to use both hands and are more sensitive to perceptual weight biases in both their dominant and non-dominant hand.

Another finding was that the perceptual bias was mainly found when lifting light objects. When lifting heavy objects, there was only a bias for right-handed participants who lifted the objects with the same right hand. In a previous study, it was also found that the perceptual bias was only seen for light objects, not heavy [5]. In another study, both for light and heavy objects a bias was found, but the effect seemed smaller for the heavy objects [6]. In both these studies only right-handed participants using their right hand to lift the objects were included, which was the only condition that showed an effect in the present study. Possibly the small effect seen for heavy object relates to the relative discrimination performance depending on the weight of the object. That is, the weight difference that can be perceived needs to be larger for heavy compared to light objects. A small perceptual bias would not be large enough to be discriminated and object weights would be judged similarly.

In line with this explanation, it is possible that the presence of the perceptual bias depends on the sensitivity to weight in a specific condition. For instance, weight discrimination is worse for passive perception, when objects are placed in the hand, compared to active lifting [27], which might explain why only a perceptual bias was found for active lifting in [5]. On the other hand, if the perceptual bias is related to weight discrimination, one would expect a smaller or absent bias in the non-dominant hand, because weight discrimination is worse for this hand [21], which is not the case for left-handers in the present study. In addition, the present study found a transfer of the perceptual bias only for left-handers, but there is no known evidence that left-handers are better at weight discrimination than right-handers [21-23]. Future research could further explore the relation between the perceptual weight bias and weight discrimination by testing these parameters in the same group of participants.

In accordance with previous research, the previous lifted object also affected the force scaling for the current lift [2-6]. The forces were scaled higher and the loading phase duration was shorter after lifting a heavy object compared to after lifting a light object. For the load force and loading phase duration, no effects of hand or handedness were found. In contrast, for the grip force rate, there were some differences dependent on the hand used or the handedness of the participant. Specifically, when using different hands, the transfer of information to the left hand seemed to be impaired. This effect was seen both in left-handed and in right-handed participants (although only for light objects in the latter group), indicating that this effect was independent of hand dominance. The difference between the used hands could suggest that information about weight used for force scaling is lateralized in the left hemisphere, which impaired a transfer of information to the right hemisphere to be used for the left hand. The lateralisation in the left hemisphere corresponds to previous studies that only found involvement of the left supplementary motor area [28] and left anterior interparietal area [29] in force control. In contrast, some previous studies did not find a difference in direction of transfer of force scaling [9, 12] or showed differences in load force, but not grip force [11], in right-handed participants. Furthermore, the finding that the transfer depends on the used hand, independent of handedness, also contradicts previous studies on motor learning with visuomotor adaptation and motor sequence learning tasks that found that transfer of learning depended on hand dominance [20, 26]. The dynamic dominance hypothesis [30, 31] suggests that the dominant and non-dominant hand have specific roles in motor control and predicts more feedforward control in the dominant hemisphere. However, the current results do not support a dependence of hand dominance. All in all, these contradictory results demand more research on the role of handedness and used hand in anticipatory force scaling for object lifting and the transfer of sensory information.

While both for perceptual estimation and anticipatory force scaling a transfer of weight information was found between hands, this depended differently on the used hand and handedness of the participant. Furthermore, although sensorimotor memory effects in peak grip force and peak load force rate were significant predictors of perceptual biases, no significant correlations were found when looking at conditions separately. Previous studies also found conflicting results concerning the correlation between force and perceptual parameters [5, 6]. van Polanen and Davare (5) proposed that corrections made to the planned forces altered the perceived weight, since the perceptual bias was only found when participants actively lifted the object, not when it was placed on their hand. However, the present results indicate that when right-handers are using different hands, these corrections to force scaling still take place but no difference in weight is perceived. While force parameters and perceptual estimates were still related on a trial-by-trial basis, the differences in findings on the transfer of information questions the hypothesis that these processes stem from the same mechanism. Weight information might be stored in different ways for perception and action or the information is transferred differently between hemispheres.

In conclusion, the present findings indicate that the sensorimotor memory from lifting an object can be transferred between hands, both in right- and left-handed participants. In contrast, a perceptual bias from lifting a previous object depended on handedness: a bias was only seen when using the same hand in right-handed participants, but could be transferred between hands in left-handed participants. In addition, in left-handers the bias was seen when using both hands, while it was only present when using the dominant hand for right-handers. These results provide insight into how information about object weight is stored and transferred between hemispheres to be used for lifting objects and perceiving their weight. These processes seem to depend differently on the used hand and handedness for perception and action.

## ACKNOWLEDGEMENTS

The author would like to thank Félice Monsieur for performing the data collection with the right-handers, and Joren Moors, Lies Vercauteren, Elise Duthoo and William De Smedt for data collection with the left-handers.

